# A modified fluctuation test for elucidating drug resistance in microbial and cancer cells

**DOI:** 10.1101/2020.11.18.389320

**Authors:** Pavol Bokes, Abhyudai Singh

## Abstract

Clonal populations of microbial and cancer cells are often driven into a drug-tolerant persister state in response to drug therapy, and these persisters can subsequently adapt to the new drug environment via genetic and epigenetic mechanisms. Estimating the frequency with which drug-tolerance states arise, and its transition to drug-resistance, is critical for designing efficient treatment schedules. Here we study a stochastic model of cell proliferation where drug-tolerant persister cells transform into a drug-resistant state with a certain adaptation rate, and the resistant cells can then proliferate in the presence of the drug. Assuming a random number of persisters to begin with, we derive an exact analytical expression for the statistical moments and the distribution of the total cell count (i.e., colony size) over time. Interestingly, for Poisson initial conditions the noise in the colony size (as quantified by the Fano factor) becomes independent of the initial condition and only depends on the adaptation rate. Thus, experimentally quantifying the fluctuations in the colony sizes provides an estimate of the adaptation rate, which then can be used to infer the starting persister numbers from the mean colony size. Overall, our analysis introduces a modification of the classical Luria–Delbrück experiment, also called the “Fluctuation Test”, providing a valuable tool to quantify the emergence of drug resistance in cell populations.

## I. Introduction

The Luria–Delbrück experiment, also called the “Fluctuation Test”, introduced 75 years ago, demonstrated that genetic mutations arise randomly in the absence of selection — rather than in response to selection — and led to a Nobel Prize. We start by reviewing this historic experiment, and highlight the subsequent development of mathematical theory to extract meaningful information from fluctuations in the data.

By the early 20th century it was known that bacteria can acquire resistance to infection by phages (bacterial viruses). However, it was debated whether mutations leading to resistance were directly induced by the virus (Lamarckian theory), or if they arose randomly in the population before viral infection (Darwinian theory). To discriminate between these alternative hypotheses, Luria & Delbhick designed an elegant experiment where single cells were isolated and grown into colonies. After allowing the colonies to grow for some duration, they were infected by the T1 phage, and the number of resistant bacteria were counted across colonies. If mutations are virus-induced (i.e., no genetic heritable component to resistance), then each bacterium has a small and independent probability of acquiring resistance, and the colony-to-colony fluctuations in the number of resistant cells should follow Poisson statistics. In contrast, if mutations occur randomly before viral exposure, then the number of surviving bacteria will vary considerably across colonies depending on when the mutation arose in the colony expansion (Fig. 1). The data clearly showed a non-Poissonian skewed distribution for the number of resistant bacteria validating the Darwinian theory of evolution [1].

**Fig. 1.**
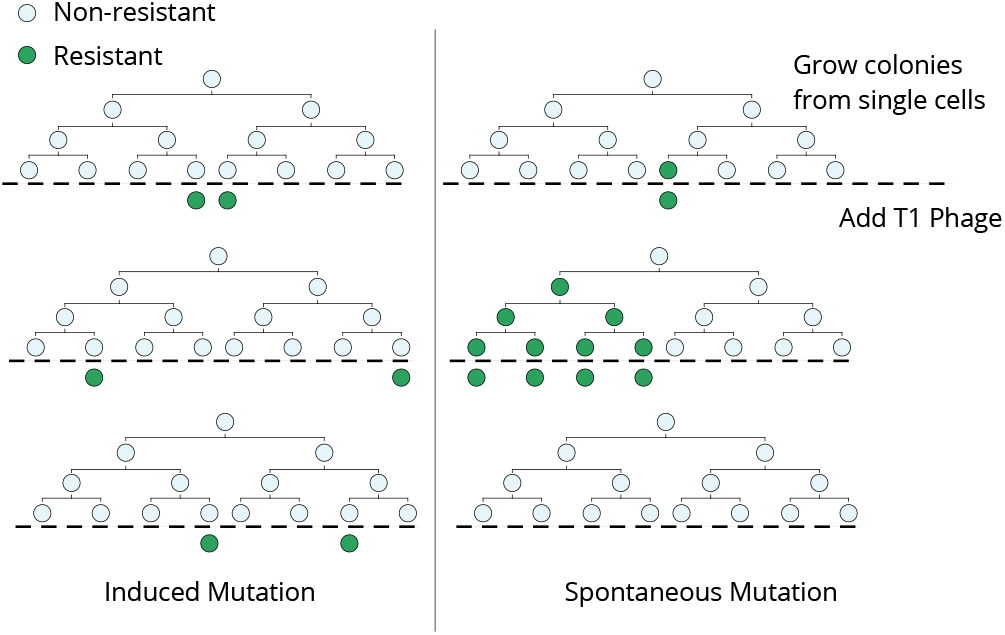
The Luria-Delbriick fluctuation test. Single cells are expanded into colonies and then infected by bacteriophage T1. If resistance mutations are virus-induced, then the number of resistant cells would follow a Poisson distribution across colonies. In contrast, if mutant cells arise spontaneously prior to viral exposure, then there will be considerable colony-to-colony fluctuations in the number of surviving cells, including “jackpot” populations where mutations happened early in the lineage expansion causing many cells to be resistant.

The Luria–Delbrück experiment, that came to be known as the “Fluctuation Test”, not only addressed a fundamental evolutionary question leading to a Noble Prize, but also laid the foundations for the field of bacterial genetics. Apart from its biological significance, the fluctuation test exemplifies the usage of stochastic analysis for uncovering hidden processes even though the underlying cell states may not be directly observable. Publication of the Luria–Delbrück article in 1943 catalyzed rich theoretical work deriving probability distributions for the number of resistant cells based on different biological assumptions [2], [3], [4], [5], and led to statistical methods for estimating mutation rates from fluctuations in the data [6], [7], [8], [9]. We refer the reader to [10] for an excellent review of mathematical developments related to the Luria–Delbrück experiment.

The fluctuation test was recently used to study cancer drug resistance [11], [12]. Individual melanoma cells were isolated from a clonal cell population by singe-cell FACS sorting, and then grown into colonies. After allowing single cells to expand for a few weeks, the colonies were treated with a targeted cancer drug, vemurafenib. Intriguingly, the colony-to-colony fluctuations in the number of surviving cells was significantly larger than a Poisson distribution, but an order of magnitude smaller than what is predicted by the mutation model (Fig. 2). Subsequent analysis showed that stochastic expression of several resistant markers drives individual cells to randomly switch between non-resistant and drug-tolerant states prior to drug exposure, and the latter state can transform into a stably-resistant state in the presence of the drug [11]. Motivated by this work we consider a system where a random number of drug-tolerant cells survive drug exposure, and then irreversibly transform to a drugresistant state with a certain adaptation rate. Our analysis shows that colony-to-colony fluctuations in the total cell count measured at single time point post drug exposure can be used to infer both the initial number of tolerant cells and their adaptation rate.

**Fig. 2.**
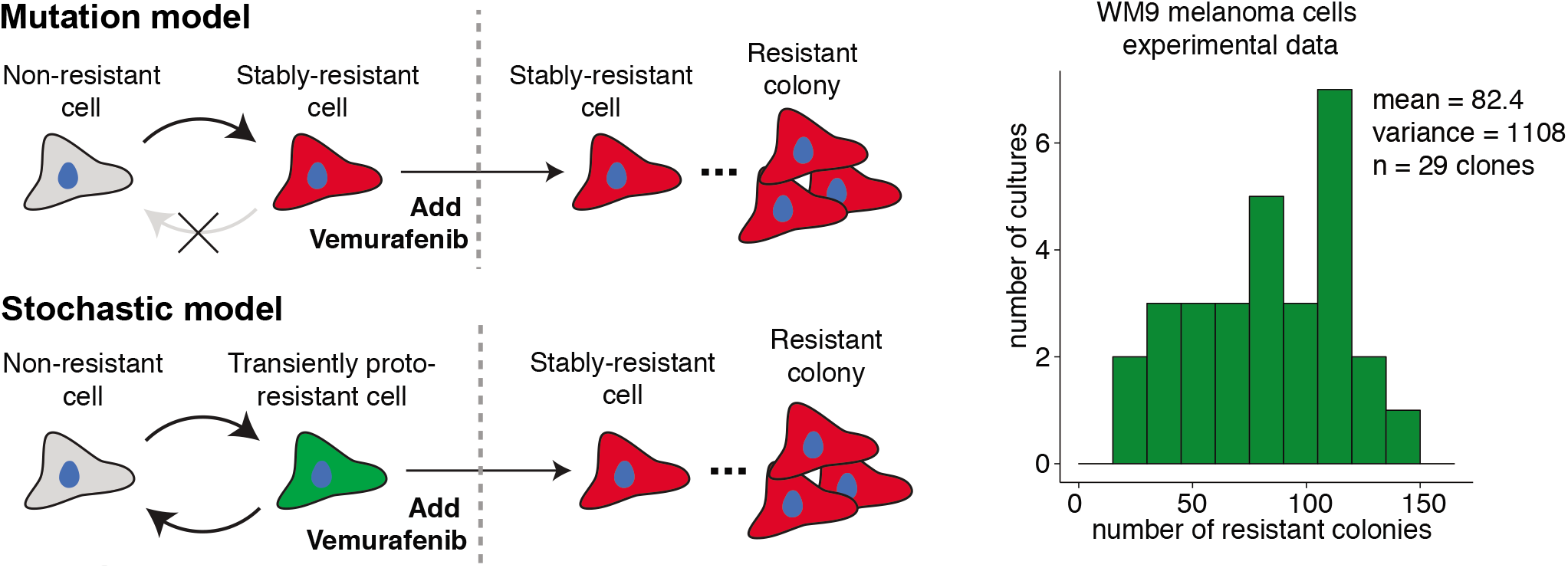
Stochastic phenotypic switching underlying cancer drug resistance. Two models for cancer drug resistance: random mutations lead to stably-resistant cells within the population prior to drug exposure (mutation model); stochastic switching between non-resistant and transiently-resistant states prior to drug exposure randomly primes a small subpopulation of cells to be drug tolerant (stochastic model). Upon drug exposure, the drug-tolerant cells survive with high probability, and transform into a stably-resistant cell. The Luria-Delbrück experiment with melanoma cells revealed non-Poissonian fluctuations in the number of surviving cells across colonies. The extent of fluctuations was significantly smaller than what is expected in the mutation model, but consistent with the stochastic model [11].

This paper is organized as follows. Section II introduces the model and its master equation. Statistical moments of the cell population size are derived in Section III. The joint distribution of the number of tolerant and resistant cells is characterized in terms of its generating function in Section IV. The solution starting from one non-resistant cell is interpreted as an exponentially delayed Yule growth process in Section V. The generating function for the total population size (tolerant plus resistant cells) is given in Section VI. Section VII provides a recursive formula for the probability mass function of the total population size for a Poisson initial condition. The paper is concluded in Section VIII.

## II. Model formulation

The number of drug-tolerant (*x*(*t*)) and drug-resistant (*y*(*t*)) cells are modelled by a bivariate continuous-time Markov chain with discrete state space {0,1,…} × {0,1,…}. The joint probability mass function *p* = *p*(*x, y, t*) of *x*(*t*) and *y*(*t*) satisfies the forward Chapman–Kolmogorov equation [13]

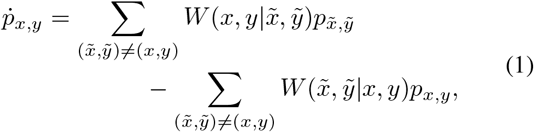

where the dot indicates the time derivative and 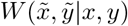 determines the transition rates in the Markov chain. They are of the following kinds:

Adaptation

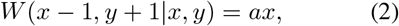

in which *a* is the adaptation rate per individual cell; Proliferation

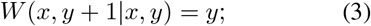

the cell doubling rate is set to one without loss of generality.

With choices (2)–(3), the forward Chapman–Kolmogorov equation (1) becomes

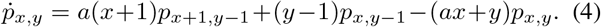

At the initial time *t* = 0, we impose an initial condition on (4) of the form

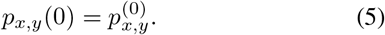

Specifically, we will focus on a situation when there are initially no drug-resistant cells and a Poisson-distributed number of drug-tolerant cells, i.e.

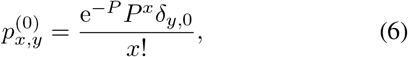

where *P* is the initial mean cell count. The symbol *δ_i,j_*, where *i* and *j* are discrete indices, represents here and below the Kronecker delta symbol, which is one if *i* = *j* and zero otherwise. Sample paths of total cell count (both drug-tolerant and drug-resistant cells) are shown in Fig. 3.

**Fig. 3.**
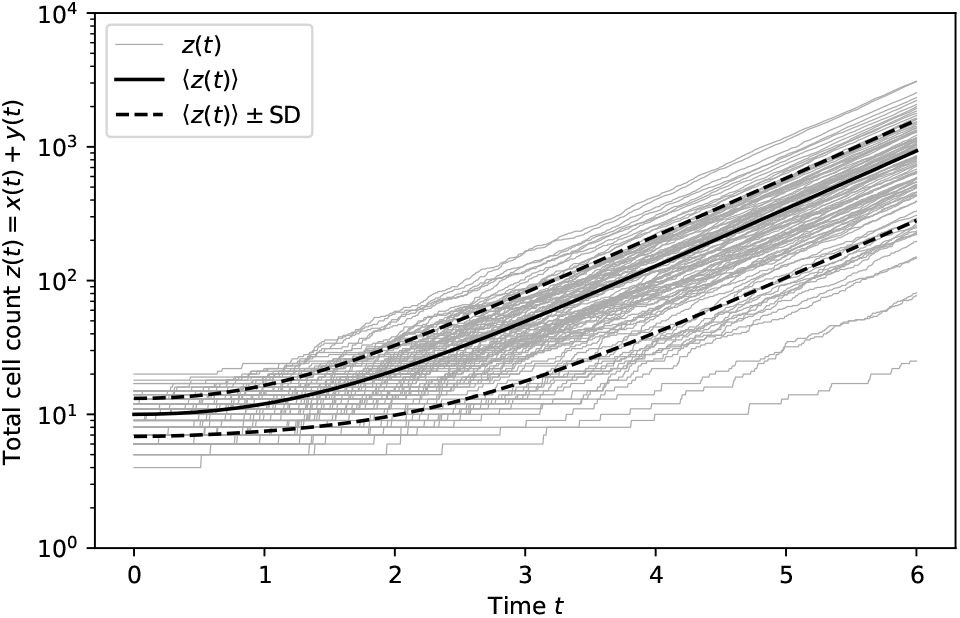
One hundred realizations of total cell count *z*(*t*) = *x*(*t*)+*y*(*t*) (gray paths) are compared to the mean value (black solid curve) and confidence limits (dashed gray curves). Initially there are only drug-tolerant cells, and their number is drawn from the Poisson distribution with mean *P* = 10. The rate of adaptation to the drug is set to *a* = 0.3 (in the units of cell doubling rate).

### III. Moments

For nonnegative integers *i* and *j* we define the joint moments of the cell counts by

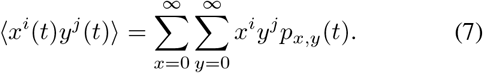

We will focus specifically on the first- and second-order moments, i.e. we take *i*, *j* ≥ 0, *i* + *j* ≤ 2.

Differential equations for the moments (7) are obtained by multiplying the master equation (4) by the monomial *x^i^y^j^* and summing over all states *x,y* ∈ {0,1…} × {0,1,…}. For the first-order moments doing so yields

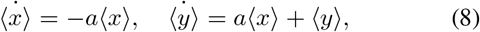

and for the second-order ones we get

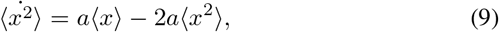

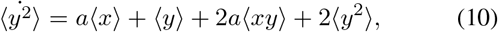

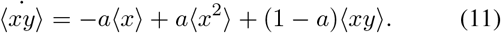

Thanks to the linear dependence of the transition rates (2)–(3) on the system state, the moment equations (8)–(11) are closed to the given (here second) order [14], [15], [16].

The moments of the initial distribution (6) are given by

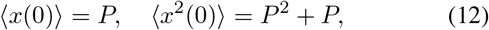

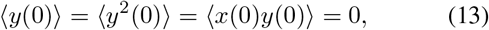

and provide the initial condition for the moment equations (8)–(11).

In addition to the partial cell counts, we are particularly interested in the total cell count, which is defined by

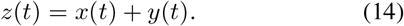

Equations

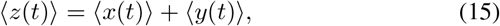

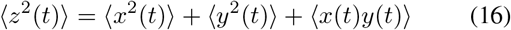

express the moments of the total cell count (14) in terms of the marginal moments (7).

Integrating the linear system of ordinary-differential equations (8)–(11) subject to the initial condition (12)–(13), and then inserting the result into (15) and (16), yields

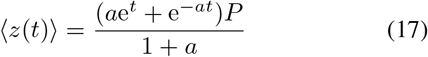

for the mean cell count and

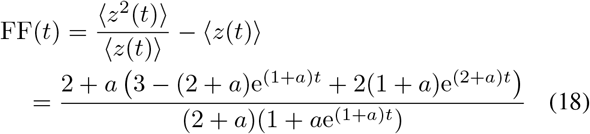

for the Fano factor (the variance-to-mean ratio). The Fano factor increases with time, exhibiting exponential growth 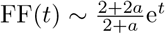 as *t* → ∞ (cf. Fig 4, top). Increasing adaptation rate increases the Fano factor. Combining (17) and (18) we obtain other useful measures of variability, namely the (squared) standard deviation SD^2^(*t*) = FF(*t*)〈z(*t*)〉 and the (squared) coefficient of variation CV^2^(*t*) = FF(*t*)/〈z(*t*)〉. The squared coefficient of variation increases with time from the Poissonian value CV^2^(0) = *P*^-1^, and plateaus at CV^2^(∞) = lim *t* → ∞FF(*t*)/〈*z*(*t*)〉 = 2(1+*a*)^2^/*a*(2+*a*)*P*. For *a* ≫ 1 (fast adaptation), CV^2^(*t*) plateaus at twice its initial value; for *a* ≪ 1 (slow adaptation), the plateau value becomes infinitely large, and the increase towards the plateau is highly sigmoid. Thus, low adaptation rates lead to small values of the coefficient of variation initially but large values at later times (cf. Fig 4, bottom).

**Fig. 4.**
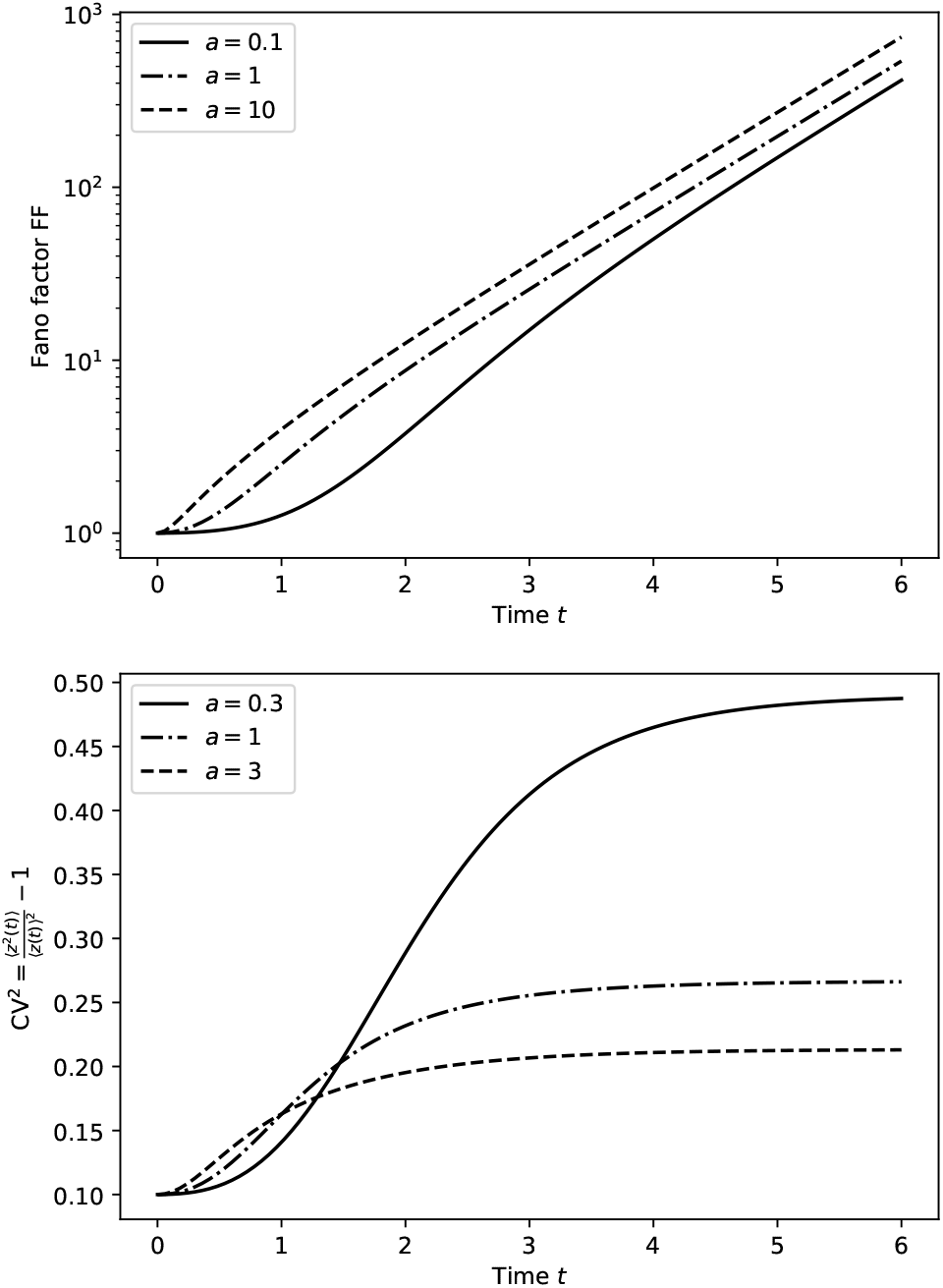
The Fano factor (upper panel) and the squared coefficient of variation (lower panel) of the total cell count as functions of time *t* for selected values (detailed inset) of the adaptation rate *a*. The initial population is drug-tolerant and its size is drawn from the Poisson distribution with mean *P* = 10.

In the next section, we go beyond the mean and variance and characterize the underlying distribution in terms of the generating function of the probability mass function.

### IV. Generating function

The joint generating function of cell counts *x*(*t*) and *y*(*t*) is defined by

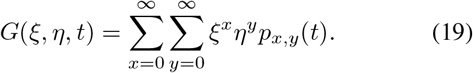

Multiplying the forward Chapman–Kolmogorov equation (4) by *ξ^x^η^y^* and summing over all states *x,y* ∈ {0,1…} × {0,1,…}, we derive a linear first-order partial differential equation for the generating function,

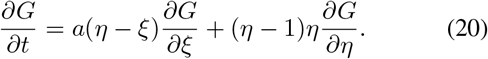

Equation (20) is subject to an initial condition that is given by the generating function of the initial distribution (5), i.e.

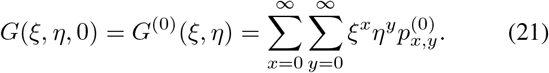

Below, we solve the initial-value problem (20)–(21) using the method of characteristics.

The characteristic system associated with the first-order partial differential equation (20) reads

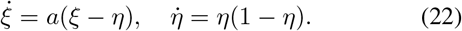

We consider a characteristic curve that passes through a point (*ξ*_0_, *η*_0_, *t*_0_) in the domain of the generating function (19), i.e. we impose a (terminal) condition

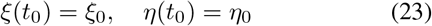

on the characteristic system (22).

Solving the logistic equation for *η* = *η*(*t*) in (22) by separating the variables *η* and *t* yields

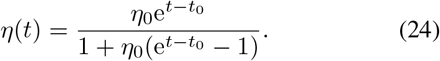

Solving the inhomogeneous linear equation for *ξ* = *ξ*(*t*) in (22) by variation of constants gives

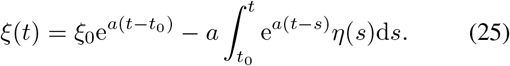

Substituting *s* = *t*_0_ – *τ* in (25) gives

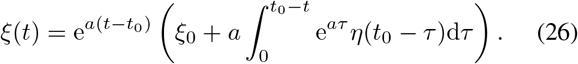

Inserting (24) into (26), we obtain

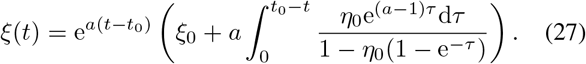

The solution of the partial differential equation (20) is constant on a characteristic curve, so that

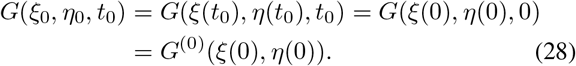

Inserting (24) and (27) with *t* = 0 into (28), and then dropping, for convenience, the subscript zero from the independent variables *ξ*_0_, *η*_0_, and *t*_0_, we arrive at

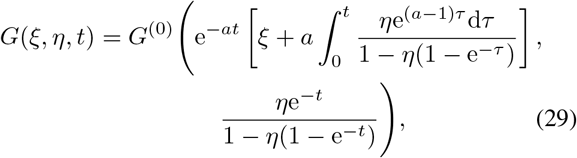

which gives the general solution to the initial value problem (20)–(21).

### V. Probabilistic interpretation

Here we supplement the formal solution given in the previous section by a probabilistic interpretation. The solution becomes tractable for simple initial conditions, in particular if initially there is (a) a single drug-resistant cell or (b) a single drug-tolerant cell. Below we show that in these two scenarios the formal solution corresponds to the Yule and the exponentially delayed Yule process, respectively.

(a) The initial presence of a single drug-resistant cell amounts to the initial distribution

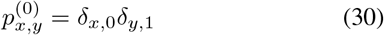

with generating function

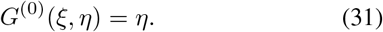

Substituting (31) into the solution formula (29) gives

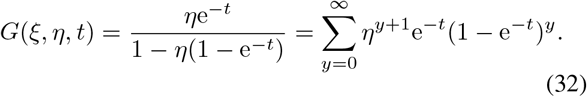

Collecting the coefficients at powers of *η*, we conclude that the number of *y* at time *t* is geometrically distributed with probability mass function *q*(1 – *q*)^*y*−1^, where *y* ≥ 1 and *q* = *e*^−1^. This is a well-known solution to the Yule process [17].

(b) The initial presence of a single drug-tolerant cell corresponds to probability mass function

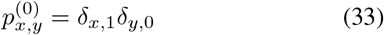

with generating function

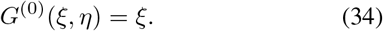

Inserting (34) into the solution formula (29), we find

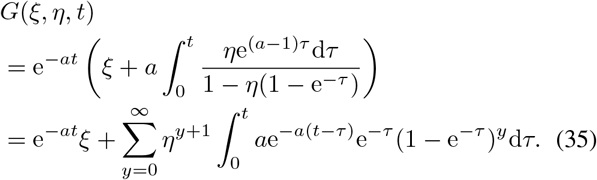

The first term in (35) pertains to the possibility that the initial cell does not adapt and remains tolerant to the drug. The probability of non-adaptation decreases exponentially in time with rate a. The remaining terms are indicative of a Yule type growth following an exponentially distributed lag. Mathematically, the delay is implemented by convolving the Yule geometric distribution with the probability density function of the exponential distribution.

### VI. Total distribution

By the total distribution we understand the distribution of the total count of cells *z*(*t*) = *x*(*t*) + *y*(*t*) (both tolerant and resistant to the drug). The total distribution can be expressed in terms of the joint probability distribution as

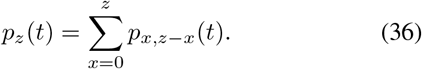

The total generating function (the generating function of the total distribution) is obtained by evaluating the joint generating function (19) along the diagonal,

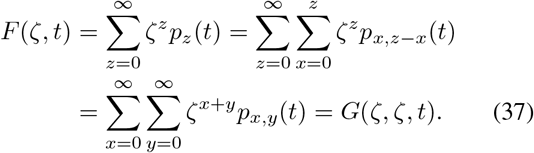

Inserting *ξ* = *η* = *ζ* into the solution formula (29), we find

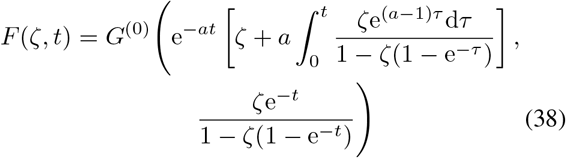

for the total generating function at time *t*.

### VII. Poissonian initial condition

Here we provide an iterative formula for the total distribution for the Poissonian initial population of tolerant cells. The method is based on an expansion of the total generating function and has previously been used in studying generalized Poisson distributions [18] and has also been applied in stochastic gene expression [19].

The generating function of the Poisson distribution (6) of initial drug-tolerant cell population is given by

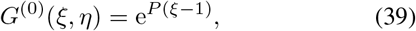

with which the total generating function formula (38) becomes

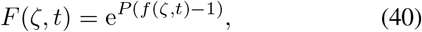

where

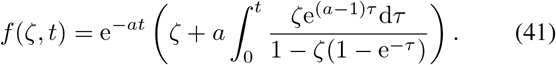

Expanding (41) in powers of *ζ*, and then substituting 1 – *e*^-*τ*^ = *u* in the integral(s), yields

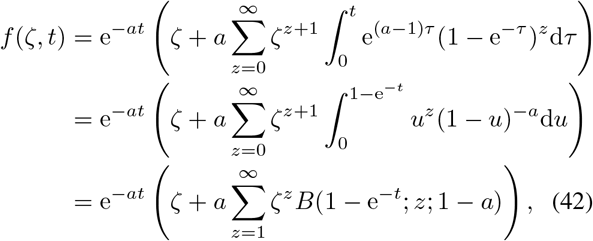

where

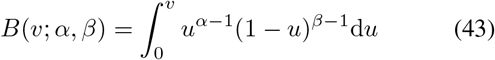

is the incomplete beta function [20].

The total probability mass function can be expressed in terms of its generating function as

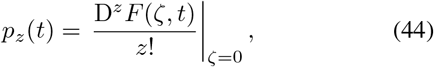

where D represents the differential operator d/dζ. For *z* = 0, (44) trivially gives the Poissonian probability of finding no cells in the initial population

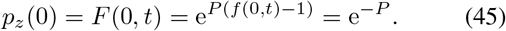

In order to find the probabilities of having positive numbers of cells, we differentiate (40) to find

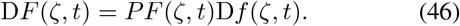

Evaluating the (*z* – 1)th derivative of (46) according to the Leibniz product rule yields

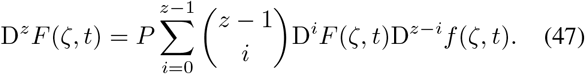

Manipulating the binomial coefficients in (47) gives

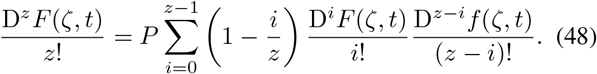

Taking *ζ* = 0 in (48) and using (44) yields

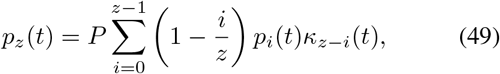

where

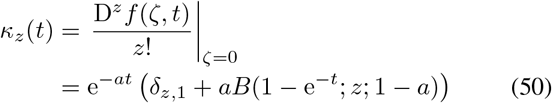

is obtained from the power-series expansion (42). Formula (49) expresses the *z*th term of the total probability mass function using the 0th, 1st, …, (*z* – 1)th terms, and can be used to evaluate the distribution recursively. The temporal evolution of the total cell count distribution is exemplified in Fig. 5.

**Fig. 5.**
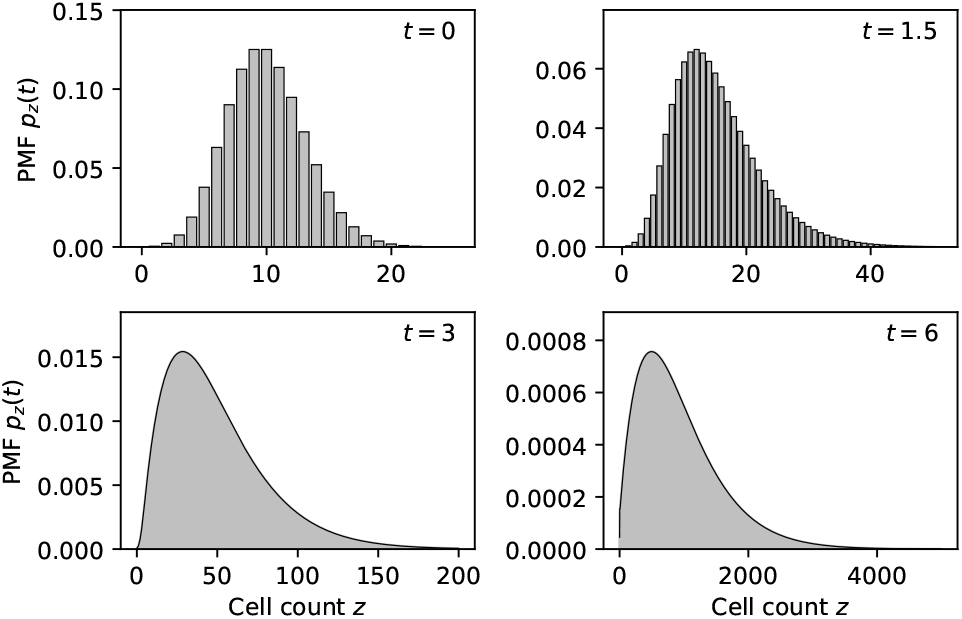
The probability mass function (45) and (49)–(50) of the total cell count for an increasing sequence of times *t.* The initial (*t* = 0) distribution is Poisson with mean *P* =10 (upper left panel). The initial population is drug-tolerant and the rate of adaptation to the drug is *a* = 0.3.

### VIII. Conclusion

In this paper, we formulated and analyzed a modification of the Luria-Delbrück experiment, in which the initially drug-tolerant cells become resistant to the treatment and begin to exponentially proliferate. We derived formulae (17)–(18) for the mean and the Fano factor and (49)–(50) for the probability mass function of the cell population size. The results are suitable for statistical inference of the adaptation rate and the initial condition.

The recursive calculation by (49)–(50) of the probability mass function remains numerically stable even at large times for large values of the cell population size. Nevertheless, we think that in future work it would be useful to complement the current discrete-distribution result by a continuous representation that would specifically apply in the large-time regime. Additionally, the current model can be extended in a number of ways, e.g. by modeling cell adaptation and/or cell cycle by a multi-step phase-type process [21], [22]. We believe that the presented analysis, and its possible generalization along the suggested lines, will contribute towards the understanding of the dynamics of drug resistance in microbial and cancer cells. Finally, the fluctuation test framework can be also used to characterize reawakening of dormant cells, such as bacterial spores, and has recently uncovered memory in the reactivation of latent HIV from human immune cells [23].

## REFERENCES

[1] S. E. Luria and M. Delbrück, “Mutations of bacteria from virus sensitivity to virus resistance,” Genetics, vol. 28, no. 6, p. 491, 1943.

[2] S. Sarkar, “Haldane’s solution of the luria-delbrück distribution,” Genetics, vol. 127, no. 2, p. 257, 1991.

[3] B. Houchmandzadeh, “General formulation of luria-delbrück distribution of the number of mutants,” Physical Review E, vol. 92, no. 1, p. 012719, 2015.

[4] C. M. Holmes, M. Ghafari, A. Abbas, V. Saravanan, and I. Nemenman, “Luria-delbrück, revisited: the classic experiment does not rule out lamarckian evolution,” Physical biology, vol. 14, no. 5, p. 055004, 2017.

[5] D. A. Kessler and H. Levine, “Large population solution of the stochastic luria-delbrück evolution model,” Proceedings of the National Academy of Sciences, vol. 110, no. 29, pp. 11682–11687, 2013.

[6] B. M. Hall, C.-X. Ma, P. Liang, and K. K. Singh, “Fluctuation analysis calculator: a web tool for the determination of mutation rate using luria-delbrück fluctuation analysis,” Bioinformatics, vol. 25, no. 12, pp. 1564–1565, 2009.

[7] A. L. Koch, “Mutation and growth rates from luria-delbrück fluctuation tests,” Mutation Research/Fundamental and Molecular Mechanisms of Mutagenesis, vol. 95, no. 2–3, pp. 129–143, 1982.

[8] M. Jones, J. Wheldrake, and A. Rogers, “Luria-delbrück fluctuation analysis: estimating the poisson parameter in a compound poisson distribution,” Computers in biology and medicine, vol. 23, no. 6, pp. 525–534, 1993.

[9] M. Kimmel and D. E. Axelrod, “Fluctuation test for two-stage mutations: application to gene amplification,” Mutation Research/Fundamental and Molecular Mechanisms of Mutagenesis, vol. 306, no. 1, pp. 45–60, 1994.

[10] Q. Zheng, “Progress of a half century in the study of the luria-delbrück distribution,” Mathematical biosciences, vol. 162, no. 1-2, pp. 1–32, 1999.

[11] S. M. Shaffer, M. C. Dunagin, S. R. Torborg, E. A. Torre, B. Emert, C. Krepler, M. Beqiri, K. Sproesser, P. A. Brafford, M. Xiao, E. Eggan, I. N. Anastopoulos, C. A. Vargas-Garcia, A. Singh, K. L. Nathanson, M. Herlyn, and A. Raj, “Rare cell variability and drug-induced reprogramming as a mode of cancer drug resistance,” Nature, vol. 546, pp. 431–435, 2017.

[12] S. M. Shaffer, B. L. Emert, R. A. Reyes Hueros, C. Cote, G. Harmange, D. L. Schaff, A. E. Sizemore, R. Gupte, E. Torre, A. Singh, D. S. Bassett, and A. Raj, “Memory sequencing reveals heritable single-cell gene expression programs associated with distinct cellular behaviors,” Cell, vol. 182, no. 4, pp. 947–959.e17, 2020.

[13] D. J. Warne, R. E. Baker, and M. J. Simpson, “Simulation and inference algorithms for stochastic biochemical reaction networks: from basic concepts to state-of-the-art,” J. Roy. Soc. Interface, vol. 16, no. 151, p. 20180943, 2019.

[14] A. Singh and J. P. Hespanha, “Stochastic analysis of gene regulatory networks using moment closure,” in 2007 American Control Conference. IEEE, 2007, pp. 1299–1304.

[15] I. Lestas, J. Paulsson, N. Ross, and G. Vinnicombe, “Noise in gene regulatory networks,” IEEE T. Circuits-I, vol. 53, no. 1, pp. 189–200, 2008.

[16] A. Singh and J. P. Hespanha, “Approximate moment dynamics for chemically reacting systems,” IEEE Transactions on Automatic Control, vol. 56, pp. 414–418, 2011.

[17] S. M. Ross, Introduction to probability models. Academic press, 2014.

[18] J. Gurland, “A generalized class of contagious distributions,” Biometrics, vol. 14, pp. 229–49, 1958.

[19] P. Bokes, J. R. King, A. T. Wood, and M. Loose, “Exact and approximate distributions of protein and mRNA levels in the low-copy regime of gene expression,” J. Math. Biol., vol. 64, pp. 829–854, 2012.

[20] M. Abramowitz and I. Stegun, Handbook of Mathematical Functions with Formulas, Graphs, and Mathematical Tables. National Bureau of Standards, Washington, D.C., 1972.

[21] M. Soltani, C. A. Vargas-Garcia, D. Antunes, and A. Singh, “Intercellular variability in protein levels from stochastic expression and noisy cell cycle processes,” lios Comput. Biol., vol. 12, no. 8, p. e1004972, 2016.

[22] C. Celik, P. Bokes, and A. Singh, “Stationary distributions and metastable behaviour for self-regulating proteins with general lifetime distributions,” in International Conference on Computational Methods in Systems Biology. Springer, 2020, pp. 27–43.

[23] Y. Lu, A. Singh, and R. D. Dar, “A transient heritable memory regulates hiv reactivation from latency,” bioRxiv, 2020. [Online]. Available: https://www.biorxiv.org/content/early/2020/07/02/2020.07.02.185215

